# *CTNS* mRNA as a potential treatment for nephropathic cystinosis

**DOI:** 10.1101/2023.04.28.538106

**Authors:** Tjessa Bondue, Sante P. Berlingerio, Florian Siegerist, Elena Sendino-Garví, Maximilian Schindler, Sara Cairoli, Bianca M. Goffredo, Fanny O. Arcolino, Jürgen Dieker, Manoe Janssen, Nicole Endlich, Roland Brock, Rik Gijsbers, Lambertus van den Heuvel, Elena Levtchenko

## Abstract

Messenger RNA (mRNA) therapies are emerging in different disease areas, but have not yet reached the kidney field. Our aim was to study the feasibility to correct the genetic defect in nephropathic cystinosis using synthetic mRNA. Cystinosis is a prototype disorder of proximal tubular dysfunction caused by mutations in the *CTNS* gene, encoding the lysosomal cystine-H^+^ symporter cystinosin, and leading to cystine accumulation in all cells of the body. The kidneys are the first and most severely affected organs, presenting glomerular and proximal tubular dysfunction. Cysteamine is the current therapeutic standard that reduces cellular cystine levels, but has many side effects and does not restore kidney function. Here, we show that synthetic mRNA is safe and effective to reintroduce functional cystinosin using lipofection in *CTNS^-/-^* kidney cells and following direct injection in *ctns^-/-^* zebrafish larvae. *CTNS* mRNA therapy results in prompt lysosomal expression of the functional protein and decreases cellular cystine accumulation for up to 14 days. In the *ctns^-/-^* zebrafish, *CTNS* mRNA therapy improves proximal tubular reabsorption, reduces proteinuria, and restores brush border expression of the multi-ligand receptor megalin. We propose that mRNA-based therapy, if sufficient kidney targeting can be achieved, may be a new approach to treat cystinosis.

**Translational statement:** Cystinosis is a systemic lysosomal storage disease caused by mutations in the *CTNS* (cystinosin) gene. It initially affects the kidneys and leads to kidney failure, if left untreated. The current standard therapy, cysteamine, is not curative and has many side-effects. Here we demonstrate the potential of mRNA-based therapy to swiftly restore cystinosin function and ameliorate the kidney phenotype. Future research will focus on mRNA delivery methods and targeting kidney cells in cystinosis rodent models.

## Introduction

Lysosomal storage disorders (LSD) are a group of over 70 monogenic diseases, affecting different lysosomal proteins and leading to lysosomal substrate accumulation.^1^ Cystinosis is an autosomal recessive LSD caused by mutations in the *CTNS* gene (chr17.13), encoding for the cystine-H^+^ symporter cystinosin. Defective cystinosin results in intra-lysosomal cystine accumulation. Recent research has demonstrated that cystinosin also regulates various metabolic and signalling pathways, including cell survival, nutrient sensing, and vesicle trafficking.^2^ The kidneys are the first and the most severely affected organs, presenting defective proximal tubular reabsorption (Renal Fanconi Syndrome, RFS) and glomerular damage leading to end-stage kidney disease (ESKD).^3,4^ Moreover, cystinosis also affects extra-renal organs, usually at adolescent or adult age.^5^

Since 1994, the standard therapy for cystinosis is the cystine-reducing agent cysteamine. While effective in prolonging kidney survival and postponing extra-renal complications, cysteamine does not reverse RFS and cannot correct the non-cystine transporting functions of cystinosin. Also, many side-effects and strict dosing schedules pose a significant burden on the patients.^6^

Preliminary results of gene therapy by transplanting *CTNS* lentiviral vector-transduced autologous hematopoietic stem cells (HSC) showed a successful reduction of tissue cystine accumulation and preserved eye, thyroid and kidney function.^7^ Nevertheless, HSC transplantation has several disadvantages, such as the risk of insertional mutagenesis and the need for leukapheresis and myeloablation.^7,8^ Messenger RNA (mRNA-)based approaches have recently gained a lot of attention during SARS-CoV2 pandemic following the extended use of mRNA vaccines.^9–11^ Additionally, mRNA therapies are currently in pre-clinical or clinical trial phase for several genetic diseases, such as cystic fibrosis and propionic acidaemia.^12–14^ mRNA acts in the cytosol, resulting in a fast translation of the therapeutic protein in dividing and non-dividing cells. Furthermore, due to the transient nature of mRNA expression, the treatment can be easily modulated or halted should adverse events occur.^15^ mRNA can be modified to reduce immunogenicity and delivery vehicles can protect it from degradation in the blood-stream or allow for organ-specific targeting, if desired. So far, mRNA-based therapies have mainly been restricted to targeting the liver and no kidney-targeted mRNA delivery vehicle has been described yet.^16,17^

In this study, we explored the potential of mRNA-based therapy to treat cystinosis in 2D and 3D cultures of *CTNS^-/-^* human kidney epithelial cells and in *ctns^-/-^* zebrafish. We choose cystinosis as a prototypical disease to evaluate the potential of this approach, as it presents with the distinct phenotype of cystine accumulation, and affects both podocytes and proximal tubular cells, offering clear readouts to evaluate the effectiveness of the treatment.

## Methods

### Cell culture

Conditionally immortalized proximal tubular epithelial cells (PTECs) and podocytes (PODOs), derived from healthy donors (WT) and cystinosis (CYS) patients (57kb deletion),^3,18–20^ were cultured in DMEM F-12 (Biowest, cat.no.L0093-500), as described before.^4,21^ The hollow fiber membrane (HFM) 3D model is representative of a functional kidney tubule and was prepared by seeding PTECs on L-DOPA and Collagen-IV coated fibers at 33°C, as described.^22^ Experiments were performed after differentiation at 37°C for 10 (PTECs) or 12 days (PODOs).

### mRNA synthesis

Human *CTNS* mRNAs (based on NM_004937.3) with C-terminal 3xHA (amino acid sequence: YPYDVPDYASYPYDVPDYAYPYDVPDYA) or mCherry-tag (KM983420.1) were synthesized by RiboPro

B.V. (Oss, The Netherlands) and de-immunized via codon optimization and dsRNA reduction. Each mRNA was equipped with translation promoting 5’ and 3’ UTRs, a 5’ anti-reverse capping analogue (ARCA), and a 150A-poly-A-tail. mRNA quality was assessed by spectrophotometry and gel electrophoresis.

### mRNA transfection

0.15µl of Lipofectamine MessengerMAX (LipoMM – Thermofisher, cat.no.LMRNA003) was mixed with 5µl of Opti-MEM medium (Gibco, cat.no.51985034) per sample and incubated for 10 minutes (min) at room temperature (RT). The solution was mixed by pipetting 5µl of Opti-MEM per 50ng of mRNA, incubated for 5min at room temperature (RT) before transfection and added to the cells (1/10 total volume).

### Transfection efficiency and protein localization

Transfected cells were fixed with 4% paraformaldehyde (Alpha Aesar, cat.no.J61899), permeabilized with 0.1% Triton X-100 (Sigma, cat.no.9002-93-1) and blocked in 2% bovine serum albumin (Sigma, cat.no.A8806), 0.2% gelatine (Sigma, cat.no.G1393) and 2% foetal bovine serum (Biowest, cat.noS181B-500) at RT. Staining for the HA-tag (mouse anti-HA) and/or lysosomal associated membrane protein 1 (rabbit anti-LAMP1)(**Supplementary Table S1**) was performed at 4°C (overnight), followed by incubation with Alexa-labelled secondary antibodies and Hoechst33342 for 45min. Transfection efficiency was quantified as proportion of AF-488 positive cells after manual counting. HFM were also stained for the 3HA-tag, F-actin and LAMP1, according to previously described protocols.^22^

### Cystine measurement in vitro

Cystine levels were measured, as described previously,^21^ in a total volume of 100µL 50mM N-ethylmaleimide (Sigma, cat.no.04259) and 50µL of 12% sulfosalicylic acid (Sigma, cat.no.S7422). Values were normalized to total protein after a Bicinchoninic acid assay (Thermofisher, cat.no. 23225).

### Fish maintenance

Zebrafish were handled and maintained in compliance with the KU Leuven animal welfare regulations (ethical approval 142/2019). Embryonic *ctns^-/-^* (*ctns* c.706C>T, p.Q236X)^23^ and *ctns^-/-^[Tg(l-fabp:DBP:eGFP)]* were used. The latter was established by crossing the *ctns^+/+^[Tg(l-fabp:DBP:eGFP)]* ^24^ fish, expressing a vitamin D-binding protein (DBP) fused with *eGFP* for detection of proteinuria, with the *ctns^-/-^* zebrafish. Larvae were subsequently screened for GFP fluorescence at 5 days post-fertilization (dpf) with the SteREO Discovery V8 microscope (Zeiss, Germany) and GFP positive larvae raised up. Genotyping was used to select *ctns^-/-^[Tg(l-fabp:DBP:eGFP)],* as described previously,^23^ and the presence of *eGFP* was confirmed by PCR.

### Microinjection of mRNA preparations in zebrafish larvae

Human mCherry-tagged *CTNS* mRNA solutions were prepared in RNAse free water (500ng/µl) with 0.05% PhenolRed (Sigma, cat.no.P0290). All mRNA injections were performed at the one-cell stage using a FemtoJet microinjector (Thermofisher) and borosilicate glass microcapillaries (World Precision Instruments, cat.no.1b100-4), prepared using a micropipette puller device (Sutter

Instruments, cat.no. Flaming/Brown p-97), or ready-made Eppendorf FemtoTips II. A tip opening diameter of ∼2µm was used and microinjections were set up with an injection volume of ∼1.7nl, resulting in a final concentration of ∼0.85ng mRNA per embryo.

### Toxicity studies in zebrafish

Embryonic health was evaluated at 5 days post-fertilization by quantification of percentage of embryonic death and morphological abnormalities.^23^

### Evaluation of CTNS-mCherry mRNA levels and protein expression in zebrafish

RNA levels after injection were determined by means of PCR at 24h, 72h and 120h. Collected embryos were sonicated in 1ml of Trizol (Thermofisher, cat.no.15596018) and phase separation was performed with chloroform (Sigma, cat.no.C2432), according to the manufacturer’s protocol. cDNA was synthesized with a mix of Oligo (dT) (Thermofisher, cat.no.18418-012), random primers (Thermofisher, cat.no.48190-011), dNTP mix (VWR, cat.no.733-1364) and SuperScript-III Reverse Transcriptase (Invitrogen, cat.no.18080093), according to the manufacturer’s protocol, and used for thermal cycling (35× 30sec at 95°C, 55°C and 72°C) with 1X GoTaq G2 Master Mixes (Promega, cat.no.M7823), 0.2µM *CTNS* and 0.2µM *bactin1* primers (**Supplementary Table S2**). Band intensity was quantified with ImageJ.^25^ Protein expression was quantified for randomly selected embryos based on fluorescence in a predetermined 200×100 pixels area in the head region.

### Cystine measurement in zebrafish

Larvae were collected in groups of 10-12 fish at 72 and 120h post-injection, washed with methylene blue-free aquarium water and homogenized in 200µl of 50mM NEM and 100µl of 12% SSA, as described.^23^ Values were normalized to total protein after a Bicinchoninic acid assay (Thermofisher, cat.no. 23225).

### Analysis of kidney function in ctns^-/-^[Tg(l-fabp:DBP:eGFP)] zebrafish

Tubular reabsorption was assessed after injection of 10kDa Alexa-647 dextran (Sigma, cat.no.T1037), as described.^23^ 16h after dextran injection, larvae were prepared for cryosectioning with the LeicaSM 200R cryomicrotome (6µm, Leica, Germany), as described before.^26^ Finally, slides were stained with Hoechst33342 and 2µg/ml CF770-conjugated wheat-germ agglutinin (biotium, cat.no.29059-1) for 10min at RT.

Proteinuria of sham and *CTNS-mCherry* mRNA injected *ctns^-/-^[Tg(l-fabp:DBP:eGFP)]* larvae was assessed by DBP-GFP fluorescence. Larvae were mounted in 96-well plates and imaged on an Acquifer Imaging Machine (Acquifer Imaging GmbH, Germany). Intravascular fluorescence intensities of the tail region were quantified with an automated FIJI script, segmenting blood vessels upon traversing blood cells in a short image sequence and measuring GFP fluorescence within the masks.

### Immunostaining of zebrafish larvae

Fish were fixed overnight in 4% PFA and washed with PBS (3x) and PBS with 0.1% Triton X-100 (Sigma, cat.no.9002-93-1). Larvae were kept overnight in 30% sucrose (Merck, cat.no.S8501), followed by embedding in gelatine (Sigma, cat.no.G7765) with 15% sucrose into a disposable mould (Electron microscopy sciences, cat.no.62352-07). Sections were made with the NX70 cryostat (4µm, Thermofisher, USA) and blocked by overnight incubation in PBS + 0.1% Triton X-100 and 5% donkey (Abcam, cat.no.ab7475) or goat serum (DAKO, cat.no.X090210-8), followed by incubation with primary antibody (overnight) and secondary Alexa-labelled antibody (2h). Sections were stained with Hoechst33342 and mounted with fluorescence mounting medium (DAKO, s3023).

### Microscopy

Images were acquired on an Operetta CLS High Content Screening Microscope (Perkin-Elmer, Germany – VIB Bio Imaging Core Leuven, Belgium), Zeiss LSM 880-Airyscan (Cell and Tissue Imaging Cluster (CIC), KU Leuven) and Eclipse CI microscope (Nikon, Japan). Zebrafish embryos were imaged with the Olympus IX71 widefield fluorescence microscope (Olympus – VIB bioimaging core Leuven, Belgium) and the SMZ18 fluorescence stereomicroscope with a P2-SHR Plan Apo 1× objective, motorized Z-drive (Nikon GMBH, Germany) and an X-Cite Xylis LED (Excelitas, Germany). 10kDa dextran images were obtained with laser scanning confocal microscopy (LC-LSM). For studying of proteinuria, zebrafish were imaged with the Acquifer Imaging Machine.

### Statistical analysis

Data were analysed using GraphPad Prism version 8.0.0 for Windows (San Diego, California USA, www.graphpad.com). Normality was assessed by a QQ-plot and equality of variances with an F-test (Brown-Forsythe). Average (±SEM) or median with 95% confidence interval (CI) [*x,x*] are presented for normal data and skewed data, respectively. Outliers were removed based on a ROUT outlier test (Q=1%). Two groups were compared with a Student’s t-test or Mann-Whitney test, multiple groups with an One-Way ANOVA or Kruskal Wallis test with Bonferroni or Dunn’s multiple testing. A double-sided p-value <0.05 was defined as being significant.

## Results

### CTNS-3HA mRNA transfection results in lysosomal cystinosin expression and reduction of cellular cystine levels after 24 hours

PTECs and PODOs from cystinosis patients were transfected with 500ng/ml of *CTNS-3HA* mRNA to study transfection efficiency and protein localization at 24 hours (24h) post-transfection. Protein expression was detected in 76.3% [64.0%,92.2%] (95% CI) of PTECs and 83.8% [77.6%,89.6%] of PODOs (**Figure 1a–b**). Co-staining for LAMP1 confirmed the lysosomal localization of cystinosin-3HA (**Figure 1c–d**). Also, in the 3D bioengineered kidney tubule, lysosomal protein expression could be confirmed post-transfection (**Figure 1e–f**).

**Figure 1:**
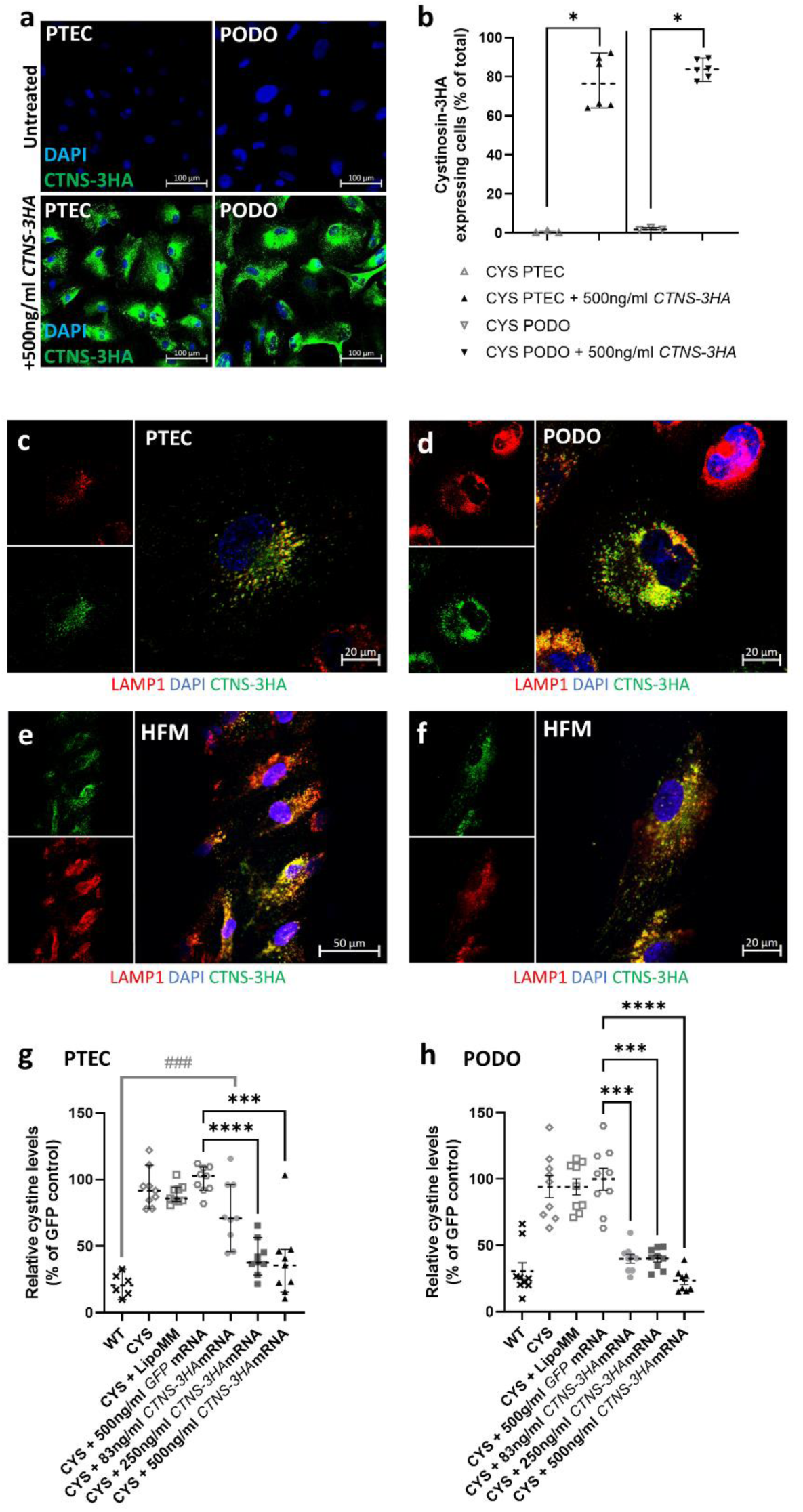
Transfection of cystinosis patient (CYS) derived proximal tubular epithelial cells (PTECs) and podocytes (PODOs) with *CTNS-3HA* mRNA results in lysosomal expression of cystinosin-3HA and reduction of lysosomal cystine accumulation at 24h post-transfection. (**a**) Cystinosin-3HA expression (green) after transfection of CYS PTECs and PODOs with 500 ng/ml *CTNS-3HA* mRNA (bottom), visualized using anti-3HA antibodies, as compared with untreated cells (top). Scale bar = 100µm. (**b**) Quantification of PTECs and PODOs expressing cystinosin-3HA in randomly acquired microscopical fields (each dot represents one observed field from 3 independent experiments). Images were obtained using Operetta CLS High Content Screening Microscope. Transfection efficiency was compared with untreated condition. Median and 95% CI are indicated. Data were analysed using individual Mann-Whitney test for each cell type. *, *P* < 0.05. (**c-f**) Lysosomal localization of cystinosin-3HA was demonstrated at 24h post-transfection by a co-staining of cystinosin-3HA (green) with LAMP1 (red) in confocal images in a 2D cell culture model (**c/d**) (scale bar = 20µm) and in 3D cell culture model of PTECs on the hollow fibre membranes (HFM – **e/f**) as functional human kidney tubules (scale bar = 50µm and 20µm, respectively). (**g/h**) Reduction of lysosomal cystine accumulation in PTECs (**g**) and PODOs (**h**) at 24h post-transfection with increasing concentrations of *CTNS-3HA* mRNA. Cellular cystine levels were analysed by LC-MS. Individual dots represent replicates from 3 experiments. *CTNS-3HA* mRNA treated cells were compared with the *GFP* control using Kruskal Wallis test (PTECs) and Welch ANOVA (PODOs)(*). A second comparison (Kruskal-Wallis) was performed with the wild type (WT) PTECs and PODOs (#). Median and 95% CI are represented for PTECs and mean with SEM for podocytes. *, *P* < 0.05; ***/###, *P* < 0.001; ****, *P* < 0.0001.

For determining whether mRNA-mediated cystinosin expression had a functional effect, cystine levels were quantified at 24h post-transfection. Three concentrations of mRNA were used to assess dose dependency. PTECs transfected with 250ng/ml or 500ng/ml of *CTNS-3HA* mRNA showed a decrease in cystine to 37.5% [28.3%, 56.4%] and 35.3% [15.5%, 47.4%] of the cystine levels observed in the CYS PTECs transfected with *GFP* mRNA, respectively. This underlines the fast-acting nature of the mRNA (**Figure 1g**). The lower concentration of *CTNS-3HA* mRNA (83ng/ml *CTNS-3HA*) showed a less pronounced decrease in cystine levels (70.8% [45.8%, 96.1%] of *GFP* control) and a smaller proportion of cells with detectable protein (**Supplementary Table S3**). Similar results were obtained in cystinotic podocytes (**Figure 1h**).

### CTNS-3HA mRNA transfection results in prolonged protein expression and cystine reduction

In the next experiments we set out to assess the duration of cystinosin expression following transfection. Cystinosin-3HA expressing PTECs were evaluated from 12h to 10 days post-transfection, with the number of cells with detectable cystinosin ranging from 84.4% [51.6%, 91.1%] at 12h to 27.1% [3.3%,46.5%] after 4 days. At 7 and 10 days, only a few cystinosin expressing PTECs could be detected (2.6% [1.0%, 5.0%] and 0.7% [0.3%, 2.5%], respectively) (**Figure 2a**). In PODOs, the fraction of positive cells ranged from 83.8% (±3.1) (±SEM) at 12h to 45.0% (±11.1) at 4 days, and a low number of cells still detectable at 7 (30.25% (±10.1)) and 10 days (10.85% (±2.0)) (**Figure 2b**).

**Figure 2:**
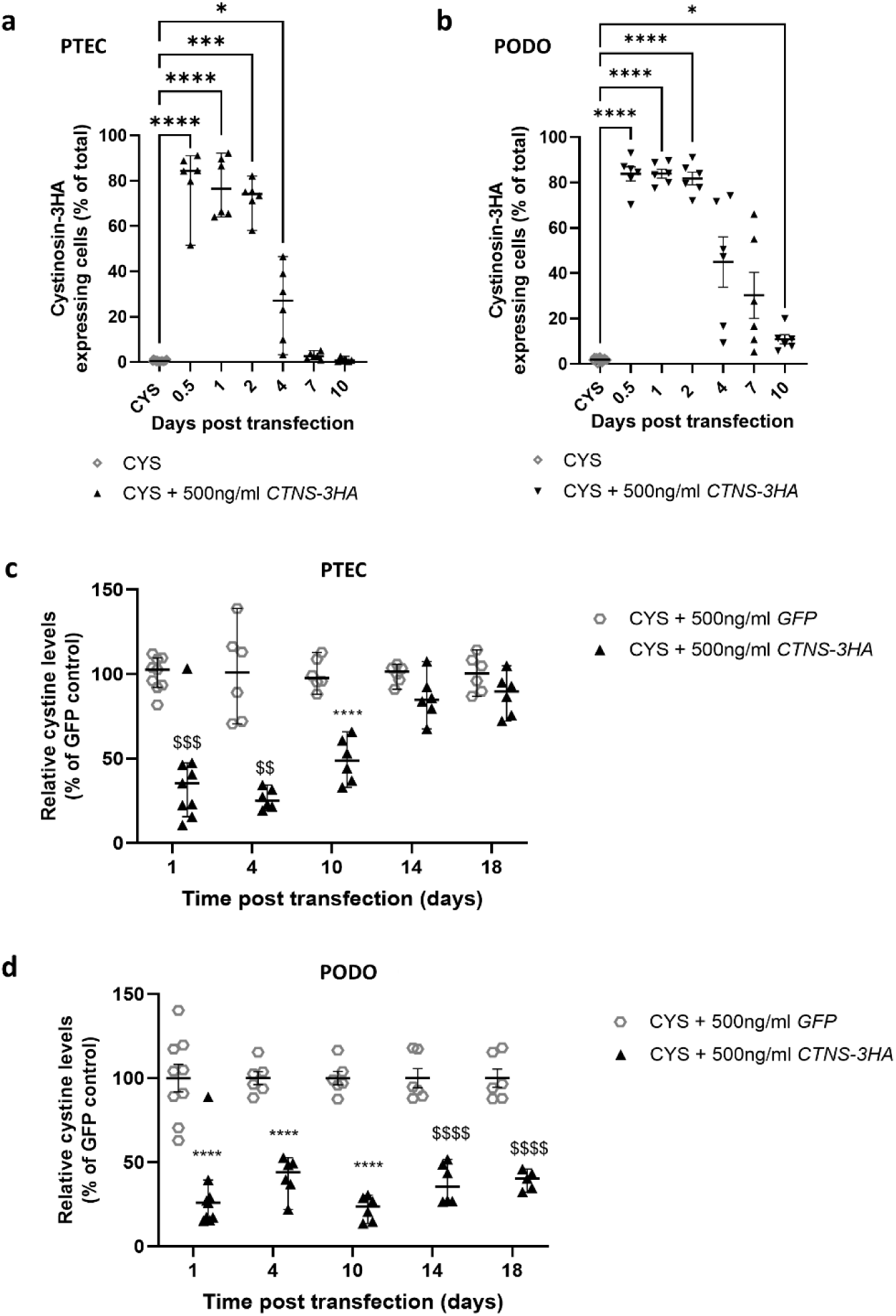
Transfection of cystinosis patient (CYS) derived proximal tubular epithelial cells (PTECs) and podocytes (PODOs) results in functional cystinosin-3HA expression for up to 4 and 10 days, and reduces cystine levels for up to 10 and 18 days, respectively. (**a/b**) Time course of cystinosin-3HA expression in PTECs (**a**) and PODOs (**b**). Images were obtained using the Operetta CLS High Content Screening Microscope and percentage of cells with detectable cystinosin-3HA expression quantified (each dot represents one observed field from 3 independent experiments). Data was analysed using a Kruskal Wallis test (PTECs) and Welch ANOVA test (PODOs). Median and 95% CI are represented for PTECs, and mean with SEM are shown for the podocytes. *, *P* < 0.05; ***, *P* < 0.001; ****, *P* < 0.0001. (**c/d**) Time course of cystine reduction in CYS PTECs (**c**) and PODOs (**d**) that were transfected with *CTNS-3HA* mRNA. Cystine levels were measured at 4, 10, 14 and 18 days post-transfection and were expressed as percentage (%) of levels measured in cells transfected with *GFP* mRNA. Each dot represents a replicate, coming from 2 independent transfections. Statistical significance was evaluated using individual tests for each time point, One-Way ANOVA (****, *P* < 0.001) and Kruskal Wallis test ($$, *P* < 0.01; $$$, *P* < 0.001; $$$$, *P* < 0.0001). Median and 95% CI are indicated for all time points.

Given the reduction in cystine levels at 24h post-transfection and detectable cystinosin protein for up to 7 days, we evaluated the long-term effect on cystine levels. Cystine levels were significantly reduced in PTECs as compared with the *GFP* mRNA control for 10 days, reaching pre-treatment levels after 14 days (**Figure 2c**). *CTNS* mRNA transfected PODOs maintained significantly lower levels for up to 18 days, with values ranging between 23.3% (± 3.0) and 40.3% [32.4%, 45.9%] of *GFP* treated cells (**Figure 2d**).

### CTNS-mCherry mRNA injection results in protein expression in ctns^-/-^ zebrafish larvae

After demonstrating the effectiveness of synthetic *CTNS* mRNA *in vitro*, we aimed to validate the mRNA-based strategy in the *ctns^-/-^* zebrafish^23^ by directly injecting human *CTNS-mCherry* mRNA into the one-cell embryo. Neither the injection procedure (sham), nor the treatment with *CTNS-mCherry* mRNA resulted in significant embryonic lethality or embryo dysmorphism, assessed as the proportion of larvae with pericardial oedema or an abnormally curved spine (**Supplementary Figure S1a-b**) at 120h post-injection (**Figure 3a–b**).

**Figure 3:**
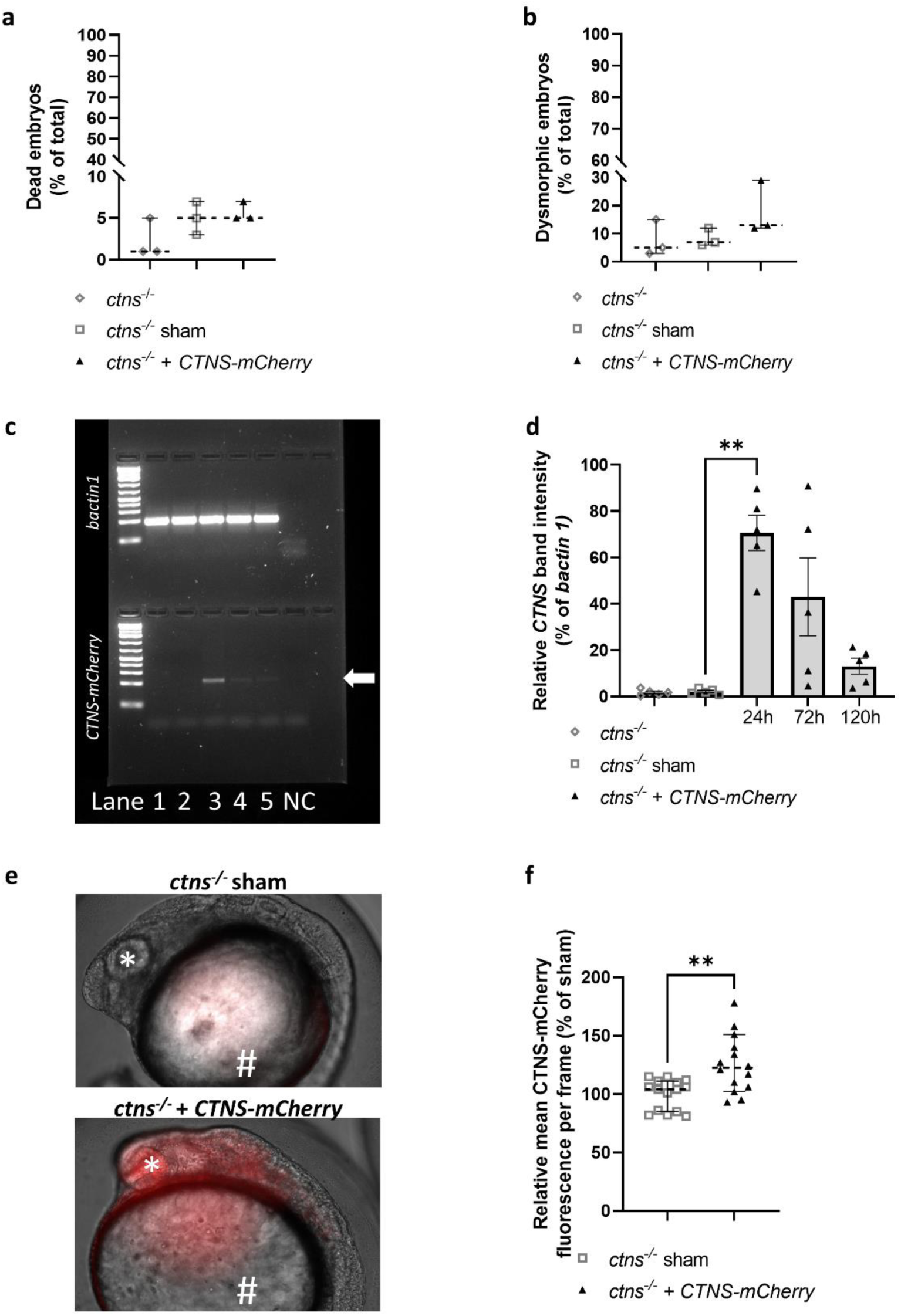
Injection of human *CTNS-mCherry* mRNA in fertilized eggs of *ctns^-/-^* zebrafish results in embryonic protein expression and does not cause toxicity. (**a/b**) The percentage (%) of dead (**a**) and dysmorphic embryos (**b**) was quantified at 5 days post-injection of human *CTNS-mCherry* (30-150 embryos per experiment, n=3). Statistical analysis was done by means of a Kruskal Wallis test. Median and 95% CI are indicated. (**c/d**) *CTNS-mCherry* mRNA levels were quantified at 24h (lane 3), 72h (lane 4) and 120h (lane 5) post-injection based on PCR band (arrow) intensity as compared with the untreated *ctns^-/-^* (lane 1) and sham treated fish (lane 2). Expression of *bactin1* mRNA was used for normalization. Per condition, 5 biological replicates are indicated (groups of 9 fish) and analysis was done with Welch ANOVA. Mean and SEM are shown. NC = negative control. **, *P* < 0.01. (**e/f**) Protein expression was quantified at 24h post-injection by live *in vivo* microscopy. Mean mCherry-fluorescence per embryo was quantified in a total of 15 embryos and normalized for the background levels observed in the sham control. Images were obtained from the larvae head region using the Olympus IX71 widefield fluorescence microscope. Head region (*) and yolk (#) are indicated. Statistical analysis was performed with a Mann-Whitney test and median and 95% CI are indicated. **, *P* < 0.01.

mRNA-levels were quantified by means of PCR band intensity and protein detection evaluated by fluorescence microscopy in the live embryo. *CTNS-mCherry* treated embryos showed high levels of *CTNS* mRNA (**Figure 3c–d**) and mCherry fluorescence (**Figure 3e–f**) at 24h (**Supplementary Figure S2a-b**). Moreover, using anti-mCherry immunostaining, we could show cystinosin expression for up to 72h (**Supplementary Figure S2c-d**).

### CTNS-mCherry mRNA injection reduces cystine accumulation in ctns^-/-^ zebrafish larvae and improves the kidney phenotype

Cystine levels were measured in zebrafish larvae at 72h and 120h post-injection and shown to decrease to 65.0% (±5.1) and 73.1% (±1.7) of levels measured in sham-treated fish, respectively (**Figure 4a**).

**Figure 4:**
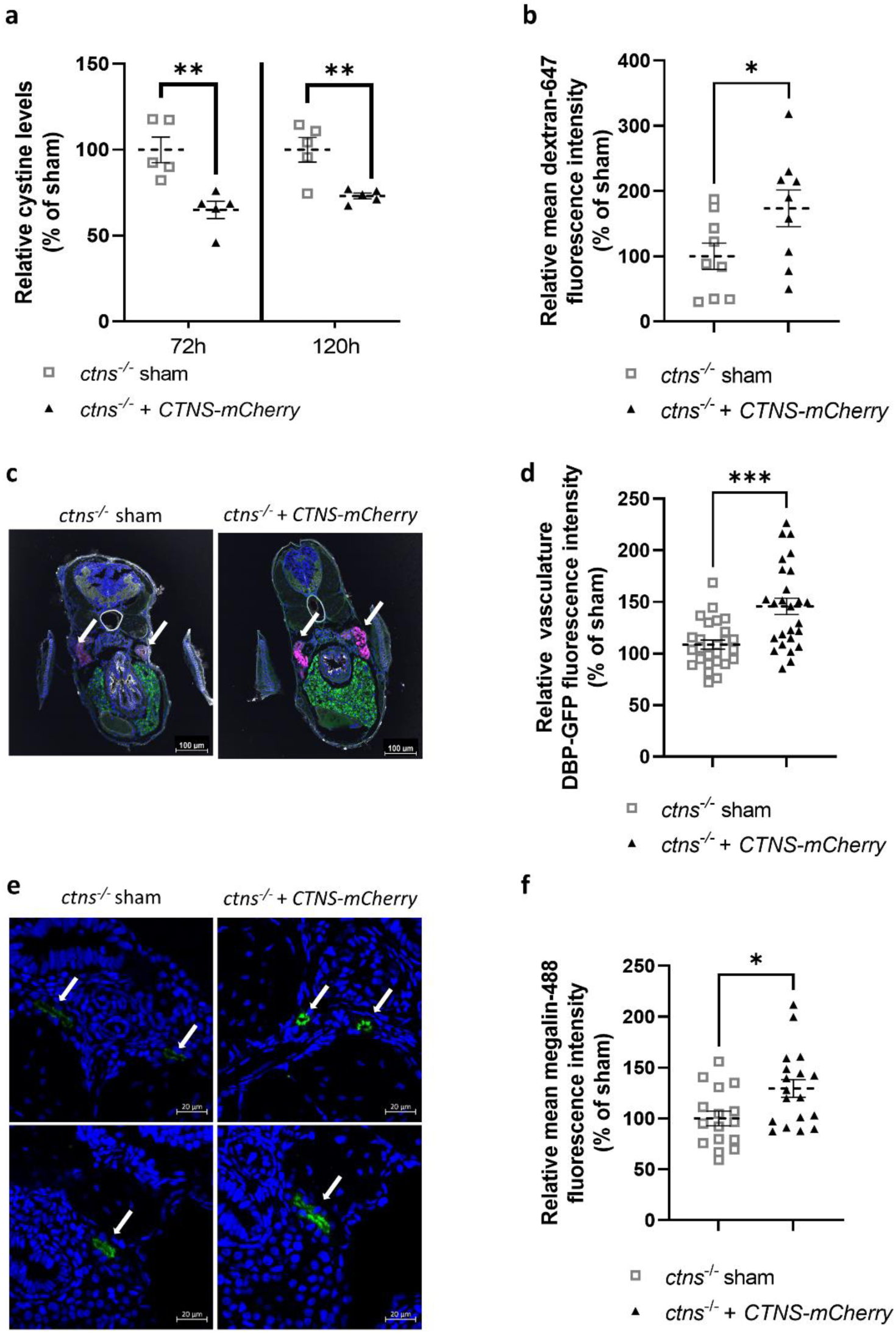
Injection of human *CTNS-mCherry* mRNA in fertilized eggs of *ctns*^-/-^ zebrafish decreases cystine accumulation, restores tubular reabsorption and alleviates glomerular proteinuria. (**a**) Injected *ctns*^-/-^ zebrafish larvae were collected in groups of 10-12 fish. Cystine levels in treated larvae were measured at 72h and 120h post-injection (n=5 groups). Statistical analysis was performed using a Student’s t-test. Mean and SEM are indicated. **, *P* < 0.01. (**b/c**) mRNA treated *ctns^-/-^[Tg(l-fabp:DBP:eGFP)]* larvae were injected with 10kDa dextran-AF647 for evaluation of low molecular weight proteinuria (LMWP) at 72h post-injection and fixed for cryosection after 16h. Sections were stained with Hoechst 33342 (blue) and wheat germ agglutinin (WGA, white), with the DBP-GFP visible in the liver (green). Tubular intensity of dextran-AF647 fluorescence (magenta) was measured and used to quantify low molecular weight protein absorption. Data were analysed with a Student’s t-test and mean with SEM is indicated (n=10 fish). Scale bar = 100µm. *, *P* < 0.05. (**d**) Glomerular proteinuria was evaluated in the *ctns^-/-^[Tg(l-fabp:DBP:eGFP)]* larvae at 120h post-injection of *CTNS-mCherry* mRNA. Images were generated with the Acquifer imaging machine. For quantification, an automatized FIJI script was used within a mask determined by segmentation of blood vessels upon traversing blood cells. Data were analysed with a Welch t-test. Mean and SEM are indicated (n=26 fish). ***, *P* < 0.001. (**e/f**) Sections of injected *ctns^-/-^* fish were prepared at 72h post-injection and stained for the megalin multi-ligand receptor on the proximal tubular brush border (green), with a DNA counterstain (blue). Confocal images were obtained and Alexa-488 intensity was measured in each tubule (n=8 fish) and showed the restauration of proximal tubular megalin expression in treated larvae. Scale bar = 20µm. Statistical significance was tested using an Student’s t-test and mean and SEM are indicated. *, *P* < 0.05.

Finally, we aimed to assess the effect of the mRNA treatment on the kidney phenotype. First, low molecular weight dextran was used to assess proximal tubular reabsorption and showed a significant improvement at 96h post-injection (**Figure 4b–c**). This was also associated with increased proximal tubular expression of megalin at 72h post-injection (**Figure 4e–f**). Subsequently, we evaluated the effect of the mRNA-based treatment on proteinuria using the *ctns^-/-^[Tg(l-fabp:DBP:eGFP)]* larvae (**Supplementary Figure S3a**).^23,24^ Here, one-cell stage mRNA injection also resulted in a detectable cystinosin expression after 24h (**Supplementary Figure S3b**) and reduced proteinuria at 120h post-injection (**Figure 4d**), as shown by an increase in embryo DBP-GFP fluorescence.

Taken together, these results demonstrate that *CTNS* mRNA-based therapy is feasible and effective, and may represent a promising approach for the treatment of cystinosis.

## Discussion

Since the discovery of the *CTNS* gene, efforts to develop a gene replacement therapy for cystinosis have been undertaken.^27^ Transplantation of *CTNS-*transduced autologous HSCs is efficient in reducing cystine accumulation and correcting different cellular pathways.^8^ However, this therapy requires leukapheresis, myeloablation and harbours other potential risks, which are not expected for mRNA-based approaches.^7,8,12^ We studied the potential of *CTNS* mRNA therapy to ameliorate cystinosis, a monogenic disease having a well-defined phenotype (cystine accumulation, and both podocyte and proximal tubular dysfunction.

A major advantage of the mRNA-based approach is the rapid expression of the protein after introduction of the mRNA into the host system.^16^ Indeed, we were able to confirm lysosomal protein expression in human cultured cells within 12h and within 6h post-injection in the zebrafish embryos. As expected, mRNA delivery resulted in a transient expression of the protein, which offers dosing flexibility and enhances safety because of the reversibility of any side effects. Moreover, mRNA therapy has no risk of insertional mutagenesis.^16^ On the other hand, the transient nature of expression can also be considered as a disadvantage, as this therapy will require repeated dosing for continuous restoration of cystinosin function. Another potential limitation of mRNA can be the initiation of an immune response due to the recognition of the mRNA by innate immune receptors (such as TLR7 and RIG-I).^16^ The immunogenicity of our mRNA was reduced through codon optimization and dsRNA removal. While the immunogenicity of mRNA was irrelevant for the cell studies, the expression of TLR7 and RIG-1 orthologues has been demonstrated in zebrafish.^28^ However, our experiments and previous studies using the injection of human mRNA in fish did not show any toxicity^29^.

The stability of mRNA can be further improved through effective 5’capping and careful selection of UTRs. As poly-A tail shortening destabilizes the RNA with every translation round, a sufficiently long (150A) poly-A tail was chosen.^30,31^ After mRNA-degradation, protein stability is the main determinant for the duration of the therapeutic effect.^12,16^ Lysosomal transporters are degraded in intraluminal vesicles during lysosomal fusion.^32^ Notably, Nevo *et al*. showed that the turnover rate of cystinosin-GFP in fibroblasts mainly depends on dilution during cell division, with the protein itself being very stable.^33^ In our study we could find cystinosin-3HA expression for up to 4 days in the kidney epithelial cells. The *CTNS* mRNA used in this study was equipped with standard translation-promoting UTRs, but the identification of UTRs tailored to the specific cell types could prolong the duration of protein expression. Cystinosin-mCherry was detectable for up 72h post-injection using immunostaining in the live zebrafish embryo. The faster decrease in detectable protein is likely caused by mRNA dilution due to the rapid cell division (from 1 to 1000 cells within 3h) during early embryo development.

Cystine accumulation, as the main hallmark of cystinosis, is the most quantifiable readout for new therapies in this disorder. Strikingly, following *CTNS* mRNA treatment we showed a decrease in cystine levels within 24h that persisted for up to 14 days (PTECs) and 18 days (PODOs). This timeframe is similar to that in mRNA-based therapies for other monogenic diseases (for example, cystic fibrosis).^13^ Notably, we observed that PTECs required higher mRNA doses (>83ng/ml) and maintained reduced cystine levels for a shorter time period as compared with the PODOs, potentially related to the higher endocytosis rate and metabolic activity of PTECs. In the zebrafish model, decreased cystine levels were observed for at least 120h. Importantly, the time course of cystine depletion exceeded that of detectable protein both *in vitro* and *in vivo*, a phenomenon that was previously shown in the study of mRNA-based therapies for Fabry disease.^34^ Therefore, we hypothesize that low levels of protein expression can still have a functional effect.

The current standard treatment of cystinosis, cysteamine, is not able to cure the RFS and can only delay the onset of ESKD. The inability of cysteamine to reverse RFS has been explained by losses of proximal tubular cells due to enhanced apoptosis.^35^ Moreover, it has been shown that cystinosin plays a direct role in regulating vesicle trafficking, autophagy and redox homeostasis in proximal tubular cells and therefore, the mere reduction of intracellular cystine is insufficient to restore proximal tubular reabsorption.^2,36–38^ Notably, Festa *et al.* found that abnormal proximal tubule dedifferentiation resulted in reduced expression of apical transporters and caused RFS in cystinosis. This phenotype was only restored after adenoviral re-expression of the *CTNS,* and not by cysteamine.^37^ In our zebrafish larvae, we demonstrated that *CTNS* mRNA addition improved proximal tubular uptake of low molecular weight dextran, a marker for proximal tubular reabsorption, and reduced overall proteinuria. In addition, we studied the expression of megalin, a multi-ligand receptor responsible for the reabsorption of low molecular weight proteins, hormones, and other polypeptides. Here, we found that treatment with *CTNS* mRNA restored receptor expression, which might indicate an overall improvement of diverse cellular pathways impaired in cystinosis proximal tubules.^39–41^

While recent studies for monogenic diseases have shown the potential of mRNA-based therapies,^12,13,42^ genetic kidney diseases lay behind.^16^ For treating the kidneys, a suitable kidney-targeting delivery vehicle that allows systemic injection of mRNA needs to be developed. Our study used a direct injection of naked mRNA into fertilized eggs of the zebrafish and provided proof-of-principle to establish effectivity of exogenously delivered mRNA. As a next step, studies in rodent models should further explore the feasibility of kidney targeting and the best administration route in more complex organisms. A suitable (kidney-targeting) delivery vehicle should protect the mRNA from degradation in the bloodstream and reduce immunogenicity by mRNA shielding.^16^ At this moment, no kidney-targeting delivery vehicles for mRNA have been described, with the targeting of the proximal tubule being the biggest challenge, since the size of the encapsulated mRNA does not allow for passage through the glomerular filtration barrier (GFM). Nevertheless, in patients with compromised podocyte function, as is the case in cystinosis or other kidney diseases, filtration of mRNA formulations might be possible. Furthermore, the basolateral uptake of 400nm nanoparticles in PTECs has been recently shown.^16,17,43^ In our study, we could already demonstrate a successful lipofectamine-mediated transfection in a 3D proximal tubule mode, mimicking basolateral delivery of mRNA.

In conclusion, our study is the first step to establishing mRNA-based therapy for cystinosis and paves the way for mRNA therapeutics in other genetic kidney diseases. Ongoing studies mainly focus on finding the appropriate delivery vehicle to allow for kidney targeting in higher organisms after systemic injection.

## Acknowledgements

We thank Marianne Klawitter (Department of Anatomy and Cell Biology, Universitätsmedizin Greifswald, Germany), Laleh Khodaparast and Ladan Khodaparast (Switch Laboratory, KU Leuven, Belgium and VIB Center for Brain & Disease Research, Leuven, Belgium), Martin Lowe (Faculty of Biology, University of Manchester, The United Kingdom), Peter de Witte and Jan Maes (Department of Pharmaceutical and Pharmacological Sciences, KU Leuven, Belgium), Kathleen Lambaerts, Frédéric Hendrickx and Kim van Kelst (Aquatic Facility, KU Leuven, Belgium) for their assistance and contribution in the zebrafish experiments. The authors gratefully acknowledge the VIB Bio Imaging Core (VIB Leuven, Belgium) for their support and assistance in this work and Pieter Vanden Berghe (Confocal Imaging Cluster (CIC), KU Leuven, Belgium) for the usage of the confocal microscope (supported by Hercules AKUL/15/37_GOH1816N and FWO G.0929.15 to Pieter Vanden Berghe, KU Leuven).

## Author contributions

TB, SPB, FS, ESG, MS, SC and BMG carried out experiments. FOA, JD, MJ, NE, RB, RG, LvdH and EL provided intellectual support, designed the study and revised the paper. RB and JD provided key reagents for the transfection. Original draft preparation was carried out by TB, SPB and FS, revision by LvdH and EL. All authors approved the final version of the manuscript.

## Legends to figures

## Supplementary materials

**Supplementary Table S1.** List of antibodies used in this study**. Supplementary Table S2.** List of primers used in this study.

**Supplementary Table S3.** Transfection efficiency in cells used for intracellular cystine measurement.

**Supplementary Figure S1.** Assessment of toxicity in *ctns*^-/-^ zebrafish. For the evaluation of toxicity following *CTNS-mCherry* mRNA injection, morphology of larvae was assessed at 120h. Two types of dysmorphism were considered, larvae presenting with pericardial oedema (a – arrow) and/or a curved spine (b). Scale bar = 500µm

**Supplementary Figure S2.** Injection of *CTNS*-*mCherry* mRNA in fertilized eggs of *ctns*^-/-^ zebrafish results in embryonic protein expression for up to 72h. (a) Cystinosin-mCherry expression was assessed at 6h, 24h, 48h, 72h, 96h and 120h post-injection in the live embryos, and mRNA treated fish were compared with the sham treated control. Images were taken using the Olympus IX71 widefield microscope. (b) Quantification of mean cystinosin-mCherry fluorescence in the head region of each fish in comparison with the sham control. Significance was tested for each time point by means of Mann-Whitney test. Median and 95%CI are shown **, *P* < 0.01. (c) In zebrafish cryosections, the mCherry-tag could also be detected by means of immunostaining using anti-mCherry antibodies in 72h old zebrafish. Images were obtained with the Nikon Eclipse CI microscope. Scale bar = 100µm. (d) Mean fluorescence per head was quantified and significance was tested compared with sham injected fish (n = 4 fish, with multiple sections analysed per fish) by means of Student’s t-test. Mean and SEM are shown. ****, *P* < 0.0001.

**Supplementary Figure S3.** *ctns^-/-^[Tg(l-fabp:DBP:eGFP)]* zebrafish present with proteinuria and express cystinosin-mCherry after injection of *CTNS-mCherry* mRNA at the one-cell stage. (a) Proteinuria was demonstrated in non-injected *ctns^-/-^[Tg(l-fabp:DBP:eGFP)]* larvae by measurement of reduced relative vasculature fluorescence intensity in comparison with the *wildtype ctns^+/+^[Tg(l-fabp:DBP:eGFP)]* larvae. Images were generated with the Acquifer imaging machine. Data were analysed with a Mann-Whitney test (n = 166 and 176 respectively), with median and 95%CI indicated. ****, *P* < 0.0001. (b) Cystinosin-mCherry expression was validated 24h after injection of *CTNS*–*mCherry* in *ctns^-/-^[Tg(l-fabp:DBP:eGFP)]* larvae. Images were obtained with the SMZ18 fluorescence stereomicroscope. Scale bar = 1mm.

