## Supplementary materials for "*CTNS* mRNA as a potential treatment for nephropathic cystinosis"

**Supplementary Table S1. List of antibodies used in this study.**

| Primary antibody | Manufacturer | Dilution |
|---|---|---|
| Anti-HA (mouse) | BioLegend, cat.no. 901515 | 1:1000 |
| Anti-LAMP1 (rabbit) | Cell signalling, cat.no. 9091S | 1:1000 |
| Anti-(zebrafish)megalin (rabbit) | homemade (Manchester, Prof. Martin Lowe) | 1:200 |
| Anti-RFP (rabbit) | Rockland, cat.no. 600-401-379 | 1:200 |
| **Secondary antibody** | **Manufacturer** | **Dilution** |
| Hoechst 33342 | Thermofisher Scientific, cat.no. 62249 | 1:1000 |
| Goat anti-Rabbit AF546 | Thermofisher Scientific, cat.no. A11035 | 1:1000 |
| Goat anti-Mouse AF488 | Abcam, cat.no. ab150113 | 1:500 |
| Donkey anti-Rabbit AF488 | Abcam, cat.no. ab150073 | 1:500 |
| Phalloidin AF647 | Thermofisher Scientific, cat.no A30107 | 1:1000 |

The anti-RFP (=red fluorescent protein) antibody was used for mCherry detection. LAMP1 = lysosomal associated membrane protein 1. AF = Alexa Fluor

**Supplementary Table S2. List of primers used in this study.**

| Gene | Forward Primer (5' -> 3') | Reverse Primer (5' -> 3') | Size of the expected band (bp) |
|---|---|---|---|
| *CTNS* (human - NM_004937.3) | CAGCGCCATTAGCATCATAAA (exon 7) | GAAACTGCTCCTTGATGTA (exon 8-9 spanning) | 212 |
| *bactin 1* (zebrafish - AF057040.1) | ACGGTCAGGTCATCACCAT | AGGGTACATGGTGGTACCTC | 191 |

**Supplementary Table S3. Transfection efficiency in cells used for intracellular cystine measurement.**

| | **Proportion of cells with detectable cystinosin-3HA protein expression*** | |
|---|---|---|
| **mRNA dose** | **PTECs** | **PODOs** |
| **83ng/ml *CTNS-3HA* mRNA** | 18% | 57% |
| **250ng/ml *CTNS-3HA* mRNA** | 63% | 67% |
| **500ng/ml *CTNS-3HA* mRNA** | 87% | 86% |

***** Representative numbers are shown, derived from a single experiment.

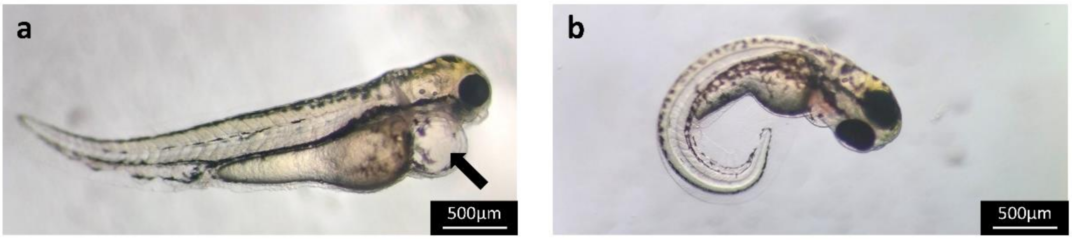


**Supplementary Figure S1. Assessment of toxicity in *ctns*$^{-/-}$ zebrafish.**

For the evaluation of toxicity following *CTNS-mCherry* mRNA injection, morphology of larvae was assessed at 120h. Two types of dysmorphism were considered, larvae presenting with pericardial oedema (**a** - arrow) and/or a curved spine (**b**). Scale bar = 500µm

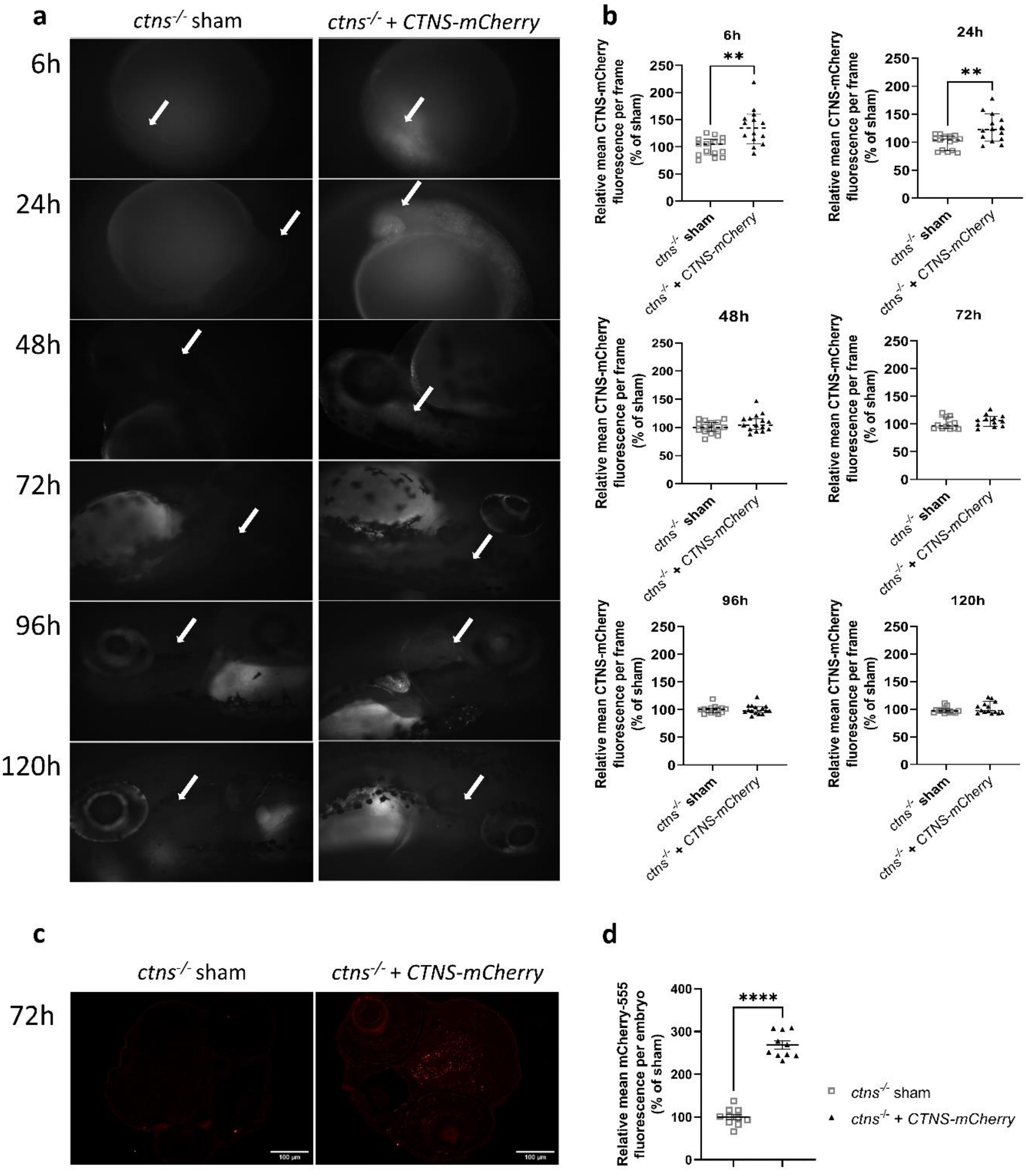


**Supplementary Figure S2. Injection of *CTNS-mCherry* mRNA in fertilized eggs of *ctns*$^{-/-}$ zebrafish results in embryonic protein expression for up to 72h.**

(**a**) Cystinosin-mCherry expression was assessed at 6h, 24h, 48h, 72h, 96h and 120h post-injection in the live embryos, and mRNA treated fish were compared with the sham treated control. Images were taken using the Olympus IX71 widefield microscope. (**b**) Quantification of mean cystinosin-mCherry fluorescence in the head region of each fish in comparison with the sham control. Significance was tested for each time point by means of Mann-Whitney test. Median and 95%CI are shown **, $P < 0.01$. (**c**) In zebrafish cryosections, the mCherry-tag could also be detected by means of immunostaining using anti-mCherry antibodies in 72h old zebrafish. Images were obtained with the Nikon Eclipse CI microscope. Scale bar = 100µm. (**d**) Mean fluorescence per head was quantified and significance was tested compared with sham injected fish (n = 4 fish, with multiple sections analysed per fish) by means of Student's t-test. Mean and SEM are shown. ****, $P < 0.0001$.

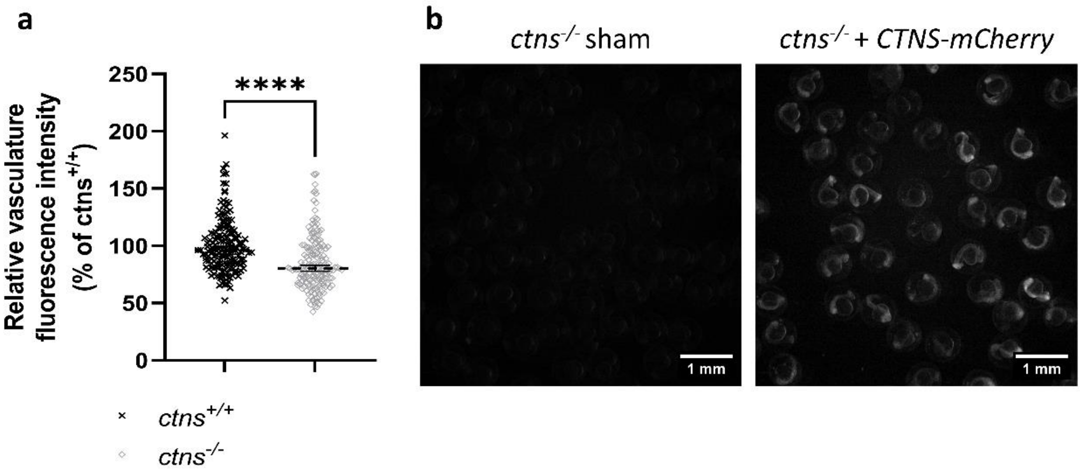


**Supplementary Figure S3.** ***ctns*$^{-/-}$*[Tg(l-fabp:DBP:eGFP)]*** **zebrafish present with proteinuria and express cystinosin-mCherry after injection of** ***CTNS-mCherry*** **mRNA at the one-cell stage.**

(**a**) Proteinuria was demonstrated in non-injected *ctns*$^{-/-}$*[Tg(l-fabp:DBP:eGFP)]* larvae by measurement of reduced relative vasculature fluorescence intensity in comparison with the *wildtype ctns*$^{+/+}$ *[Tg(l-fabp:DBP:eGFP)]* larvae. Images were generated with the Acquifer imaging machine. Data were analysed with a Mann-Whitney test (n = 166 and 176 respectively), with median and 95%CI indicated. ****, $P < 0.0001$. (**b**) Cystinosin-mCherry expression was validated 24h after injection of *CTNS-mCherry* in *ctns*$^{-/-}$*[Tg(l-fabp:DBP:eGFP)]* larvae. Images were obtained with the SMZ18 fluorescence stereomicroscope. Scale bar = 1mm.
